# How many specimens make a sufficient training set for automated 3D feature extraction?

**DOI:** 10.1101/2024.01.10.575054

**Authors:** James M. Mulqueeney, Alex Searle-Barnes, Anieke Brombacher, Marisa Sweeney, Anjali Goswami, Thomas H. G. Ezard

**Author notes:** **Author of Correspondence:** James M. Mulqueeney. **<u>Author-supplied statements</u>** Relevant information will appear here if provided. **Ethics** Does your article include research that required ethical approval or permits?: This article does not present research with ethical considerations Statement (if applicable): CUST_IF_YES_ETHICS: No data available. **Data** It is a condition of publication that data, code and materials supporting your paper are made publicly available. Does your paper present new data?: Yes. Statement (if applicable):* All computer code and final raw data used in the paper is available at https://github.com/JamesMulqueeney/Automated-3D-Feature-Extraction. All other data is available in the Dryad Repository: DOI: 10.5061/dryad.x69p8czrb. (This data is not yet made fully publically available, and which it to remain so until accepted for publication; the domains have been created and are private at this moment). ***Conflict of interest*** I/We declare we have no competing interests. Statement (if applicable):* CUST_STATE_CONFLICT: No data available.

## Abstract

Deep learning has emerged as a robust tool for automating feature extraction from 3D images, offering an efficient alternative to labour-intensive and potentially biased manual image segmentation methods. However, there has been limited exploration into the optimal training set sizes, including assessing whether artificial expansion by data augmentation can achieve consistent results in less time and how consistent these benefits are across different types of traits. In this study, we manually segmented 50 planktonic foraminifera specimens from the genus Menardella to determine the minimum number of training images required to produce accurate volumetric and shape data from internal and external structures. The results reveal unsurprisingly that deep learning models improve with a larger number of training images and that data augmentation can enhance network accuracy by up to 8.0%. Notably, predicting both volumetric and shape measurements for the internal structure poses a greater challenge compared to the external structure, due to low contrast between different materials and increased geometric complexity. These results provide novel insight into optimal training set sizes for precise image segmentation of diverse traits and highlight the potential of data augmentation for enhancing multivariate feature extraction from 3D images.

**Subject Category:** Life Sciences – Computer Science

**Subject Areas:** computational biology, Artificial Intelligence

## 1. Introduction

Three-dimensional imaging techniques, such as x-ray micro-computed tomography (micro-CT), have revolutionised the characterisation of both internal and external structures of diverse objects. With the ability to generate high-resolution images, researchers can visualise and quantify intricate 3D features with wide-ranging applications [1, 2]. However, objective extraction of these features remains a major challenge.

Currently, the prevailing approach to extracting 3D features from image data involves manual segmentation, which is a labour-intensive [3] and subjective process that lacks reproducibility [4]. This manual approach limits the study of a large number of samples and the exploration of complex hypotheses. As the acquisition of high-resolution scans has increased steadily [5, 6], there is a pressing need to enhance the efficiency of this important processing step.

Machine learning techniques, particularly deep learning, and convolutional neural networks (CNNs) offer a promising solution for automating image segmentation. These methods can potentially accelerate this processing step to deliver accurate and repeatable results, whilst being accessible to various research fields [7, 8]. However, despite their merits, a fundamental trade-off exists between the quantity of samples necessary for generating accurate neural networks and the time-consuming, subjective nature of manual segmentation for evaluation. The specific number of required samples is likely to vary across datasets and depends on the traits being extracted. Hence, there is an imperative to establish the minimum number of manually segmented specimens needed to train these neural networks and to understand how they vary in different trait extraction scenarios.

Strategies to generate training sets that reduce manual processing while maintaining performance are required. Smart interpolation, whereby pre-segmented slices and the volumetric image data are used to predict segmentation across the entire specimen, is one suggested approach [9, 10], but remains time-consuming for creating large training sets. Data augmentation, which artificially expands the size of the training set without collecting new data, can mitigate overfitting and bolster the accuracy of CNNs during training [11, 12]. Consequently, data augmentation techniques may allow smaller training sets to achieve equivalent accuracy levels of their larger counterparts.

In this study, we aim to determine the minimum number of images needed to train a neural network to produce segmentation data that is indistinguishable from manually generated data. To do so, we use a dataset of computed tomography (CT) scans of planktonic foraminifera from the genus *Menardella.* We assess the efficacy of each training set in extracting volumetric and shape data for the external calcite and internal chamber space of selected specimens (see figure 1). Additionally, we introduce a novel 3D data augmentation technique to bolster training sets by generating six different orientations of each specimen through rotation. This comparison serves to assess how data augmentation strategies can improve training sets to achieve accurate and efficient 3D feature extraction.

**Figure 1.**
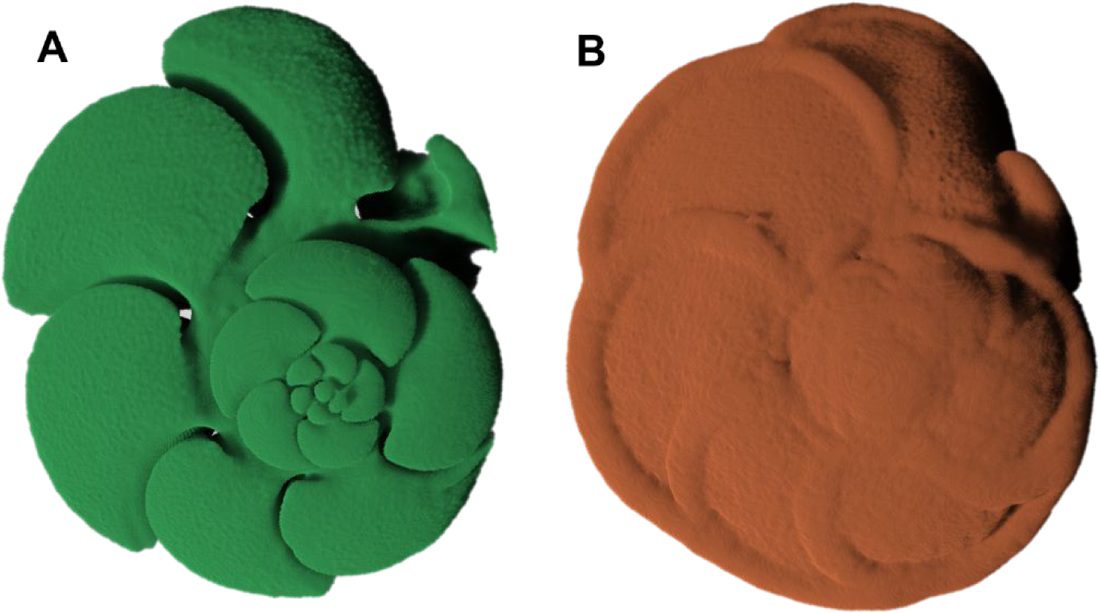
3D models of the (a) internal and (b) external structure of planktonic foraminifera generated using manual segmentation in Dragonfly v. 2021.3 (Object Research Systems, Canada).

## 2. Methods and Materials

### 2.1 Data collection

50 planktonic foraminifera, comprising 4 *Menardella menardii*, 17 *Menardella limbata*, 18 *Menardella exilis*, and 11 *Menardella pertenuis* specimens, were used in our analyses. These were collected from the Ceara Rise in the Equatorial Atlantic region at Ocean Drilling Program (ODP) Site 925, which comprised Hole 925B (4°12.248’N, 43°29.349’W), Hole 925C 20 (4°12.256’N, 43°29.349’W), and Hole 925D (4°12.260’N, 43°29.363’W). See Curry et al., [13] for more details. The samples used spanned 5.65 million years ago (Ma) to 2.85 Ma [14].

The non-destructive imaging of both internal and external structures of the foraminifera was conducted at the µ-VIS X-ray Imaging Centre, University of Southampton, UK, using a Zeiss Xradia 510 Versa X-ray tomography scanner. Employing a rotational target system, the scanner operated at a voltage of 110 kV and a power of 10 W. Projections were reconstructed using Zeiss Xradia software, resulting in 16-bit greyscale .tiff stacks characterised by a voxel size of 1.75 μm and an average dimension of 992 x 1015 pixels for each 2D slice.

### 2.2 Generation of training sets

We extracted the external calcite and internal cavity spaces from the micro-CT scans of these 50 planktonic foraminiferal specimens using manual segmentation within Dragonfly v. 2021.3 (Object Research Systems, Canada). This step took approximately 480 minutes per specimen (24,000 minutes total) and involved the manual labelling of 11,947 2D images. Segmentation data for each specimen were exported as multi-label (3 labels: external, internal, and background) 8-bit multipage .tiff stacks and paired with the original CT image data to allow for training (see figure 2).

**Figure 2.**
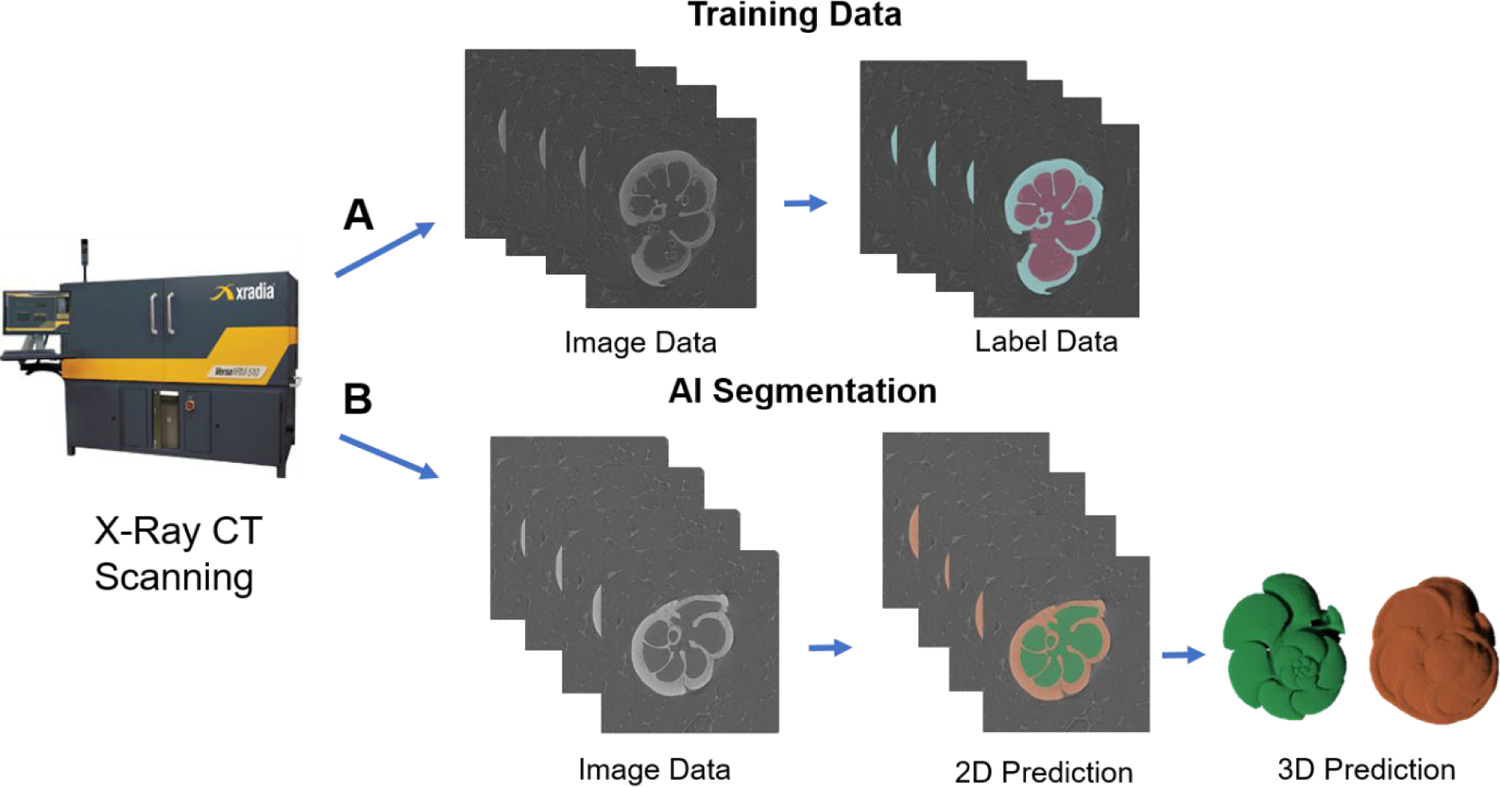
Workflow for producing training data and applying a deep convolutional neural network (CNN) to perform automated image segmentation. The workflow includes (a) the creation of training data for the input into Biomedisa and (b) an example application of the trained CNN to automate the process of generating segmentation (label) data.

The 50 specimens were categorised into three distinct groups (see supplementary table 1): 20 training image stacks, 10 validation image stacks, and 20 test image stacks. From the training image category, we generated six distinct training sets, varying in size from 1 to 20 specimens (see table 1). These were used to assess the impact of training set size on segmentation accuracy, as determined through a comparative analysis against the validation set (see Section 2.3).

**Table 1.** Design of the training sets detailing the name of the training set, the data type (original or new augmentation), the number of specimens in each training set, the number of 2D images available for training and the approximate time to perform the manual segmentation to create the training set.

| Name of Training Set | Data Type | Number of Specimens | No. of foreground Images (XY) | Approx. Duration of Segmentation (Mins) |
| --- | --- | --- | --- | --- |
| T0_AI_1_Images | Original | 1 | 241 | 480 |
| T1_AI_2_Images | Original | 2 | 390 | 960 |
| T2_AI_4_Images | Original | 4 | 1028 | 1920 |
| T3_AI_8_Images | Original | 8 | 2187 | 3840 |
| T4_AI_16_Images | Original | 16 | 3571 | 7680 |
| T5_AI_20_Images | Original | 20 | 4251 | 9600 |
| Aug_T0_AI_1_Images | Augmentation | 1 | 4480 | 480 |
| Aug_T1_AI_2_Images | Augmentation | 2 | 8812 | 960 |
| Aug_T2_AI_4_Images | Augmentation | 4 | 18120 | 1920 |
| Aug_T3_AI_8_Images | Augmentation | 8 | 36398 | 3840 |
| Aug_T4_AI_16_Images | Augmentation | 16 | 71100 | 7680 |
| Aug_T5_AI_20_Images | Augmentation | 20 | 88432 | 9600 |

From the initial six training sets, we created six additional training sets through data augmentation using the NumPy library [15] in Python. This augmentation process entailed rotating the original images five times, effectively producing six distinct 3D orientations per specimen for each of the original training sets (see figure 3). The augmented training sets comprised between 6 and 120 .tiff stacks (see table 1).

**Figure 3.**
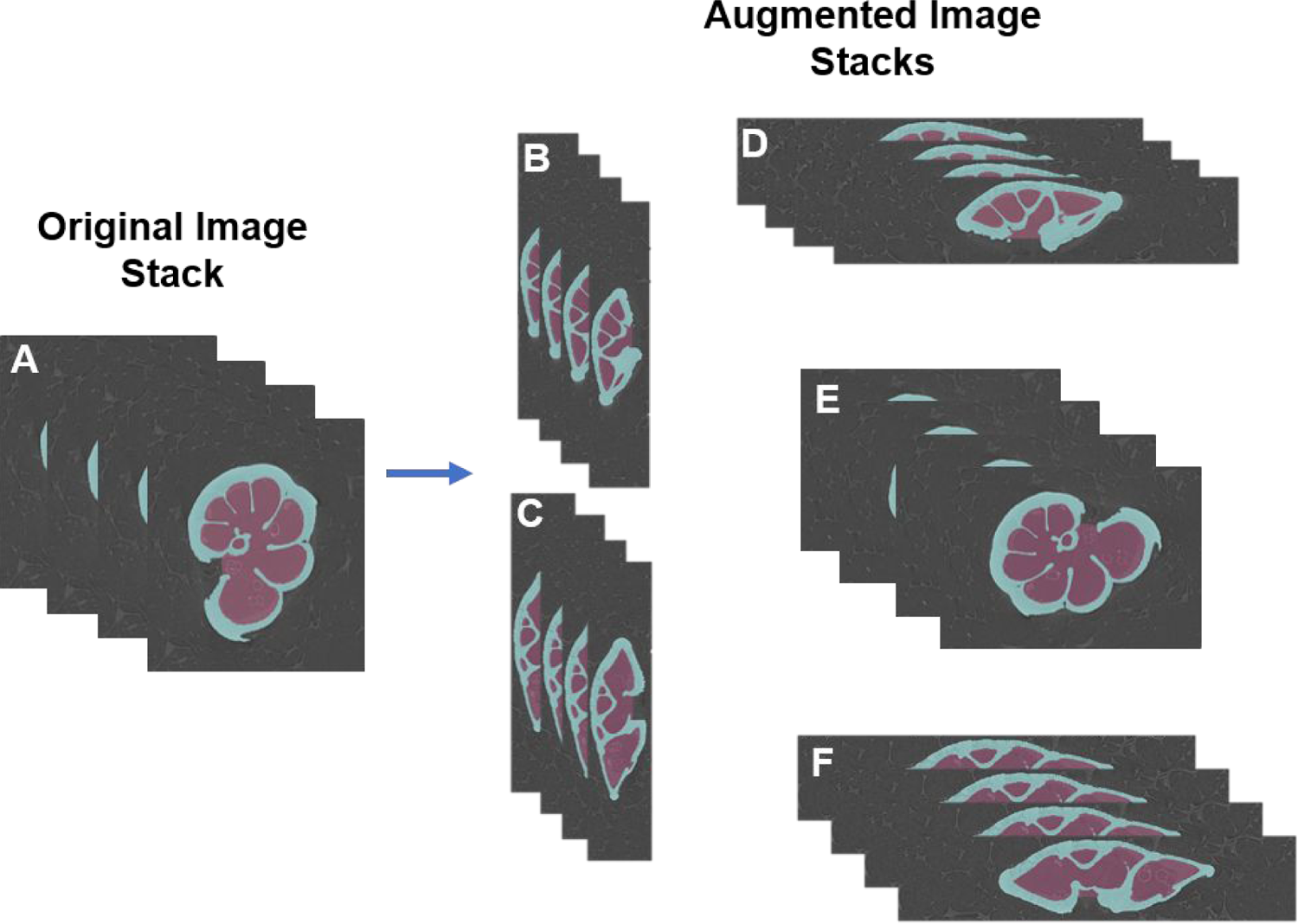
Rotation of the original image to form the new augmentation training data. The original image in an (a) xyz orientation is rotated into 5 other three-dimensional planes: (b) yzx, (c) zyx, (d) xzy, (e) yxz, and (f) zxy orientations. These are then all paired together and used in training.

### 2.3 Training the neural networks

CNNs were trained using the offline version of Biomedisa, which utilises a U-Net architecture [16] and is optimised using Keras with a TensorFlow backend. We used patches of size 64 x 64 x 64 voxels, which were then scaled to a size of 256 x 256 x 256 voxels. We trained 3 networks for each of the training sets to check the extent of stochastic variation on the results [17].

To train our models in Biomedisa, we used a stochastic gradient descent with a learning rate of 0.01, a decay of 1 × 10^-6^, momentum of 0.9, and Nesterov momentum enabled. A stride size of 32 and a batch size of 24 samples per epoch were used alongside an automated cropping feature, which has been demonstrated to enhance accuracy [18]. The training of each network was performed on a Tesla V100S-PCIE-32GB graphics card with 30989 MB of available memory. All the analyses and training procedures were conducted on the High-Performance Computing (HPC) system at the Natural History Museum, London.

To measure network accuracy, we used the Dice similarity coefficient (Dice score), a metric commonly used in used in biomedical image segmentation studies [19, 20].

The Dice score quantifies the level of overlap between two segmentations, providing a value between 0 (no overlap) and 1 (perfect match). For two segmentations, *X* and *X*^′^ consisting of n labels, the Dice score is defined as:

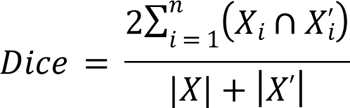

Where |*X*| and |*X*^′^| are the total number of voxels of each segmentation, respectively, and *X*_i_ is the subset of voxels of *X* with label *i*.

We conducted experiments to evaluate the potential efficiency gains of using an early stopping mechanism within Biomedisa. We opted for an early stopping criterion set at 25 epochs, which means if there is no increase in Dice score within a 25-epoch window, the optimal network is selected, and training is terminated. To gauge its impact of early stopping on network accuracy, we compared the results obtained from the original six training sets under early stopping to those obtained on a full run of 200 epochs.

### 2.4 Evaluation of feature extraction

We used the median accuracy network from each of the 12 training sets to produce segmentation data for the external and internal structures of the 20 test specimens. We then compared the volumetric and shape measurements from the manual data to those from each training set. The volumetric measurements were total volume (comprising both external and internal volumes) and percentage calcite (calculated as the ratio of external volume to internal volume, multiplied by 100).

To compare shape, mesh data for the external and internal structures was generated from the segmentation data of the 12 training sets and the manual data. Meshes were decimated to 50,000 faces and smoothed before being scaled and aligned using Python and Generalized Procrustes Surface Analysis (GPSA) [21], respectively. Shape was then analysed using the landmark-free morphometry pipeline, as outlined by Toussaint et al., [22]. We used a kernel width of 0.1mm and noise parameter of 1.0 for both the analysis of shape for both the external and internal data, using a Keops kernel (PyKeops; https://pypi.org/project/pykeops/) as it performs better with large data [22]. The analyses were run for 150 iterations, using an initial step size of 0.01. The manually generated mesh for the individual st049_bl1_fo2 was used as the atlas for both the external and internal shape comparisons.

## 3. Results

### 3.1 Early stopping vs. 200 epochs

A likelihood ratio test found no detectable difference between the observed correlation of the six original training sets under early stopping within 25 epochs and the full run of 200 epochs (F_4, 5_ = 0.0424, p > 0.05), with a strong correlation between the early stopping and full run values (see figure 4; R^2^ = 0.9997, p < 0.001). Consequently, we report results using the early stopping criterion for all subsequent tests.

**Figure 4.**
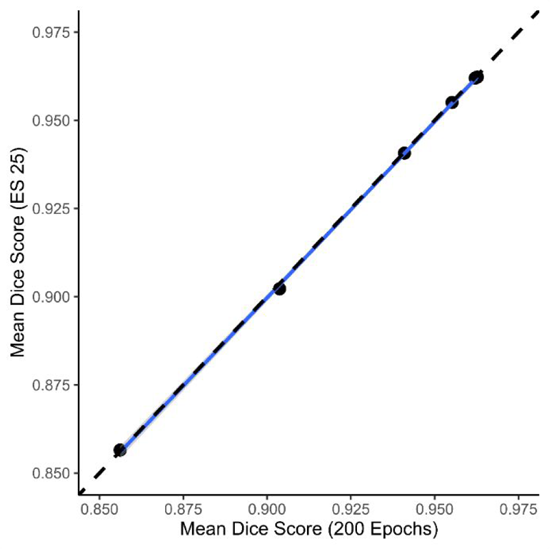
Comparison of Dice scores obtained from early stopping within 25 epochs and a full run of 200 epochs for each of the 6 original image training sets shows no statistical difference. The correlation coefficient (R^2^) between the two sets of scores displayed is 0.9998.

### 3.2 Network accuracy

The Dice scores of both the original and augmented datasets increased with the number of training images (figure 5). The improvement in accuracy from 1 specimen to 20 specimens was 12.34% and 4.67% for the original and augmented datasets, respectively. Most of this improvement occurred between training sets of 1 to 8 specimens. An Analysis of Variance (ANOVA) test showed that the differences in Dice scores varied across all training sets (F_11, 24_ = 92.84, p < 0.001). The augmented training data resulted in higher mean Dice scores compared to their original counterparts, especially at lower specimen numbers (figure 5). A generalised linear model with a quasibinomial error structure showed that increasing specimen numbers increased the Dice score for the augmented data (*β* = 0.314 on the scale of the logit link function, s.e. = 0.027, p < 0.001), but did so faster for the original data (*β* = 0.211 on the scale of the logit link function, s.e. = 0.031, p < 0.001) but from a much lower initial baseline (*β* = −0.633 on the scale of the logit link function, s.e. = 0.0059, p < 0.001). The model explains 96% of the variation in the data.

**Figure 5.**
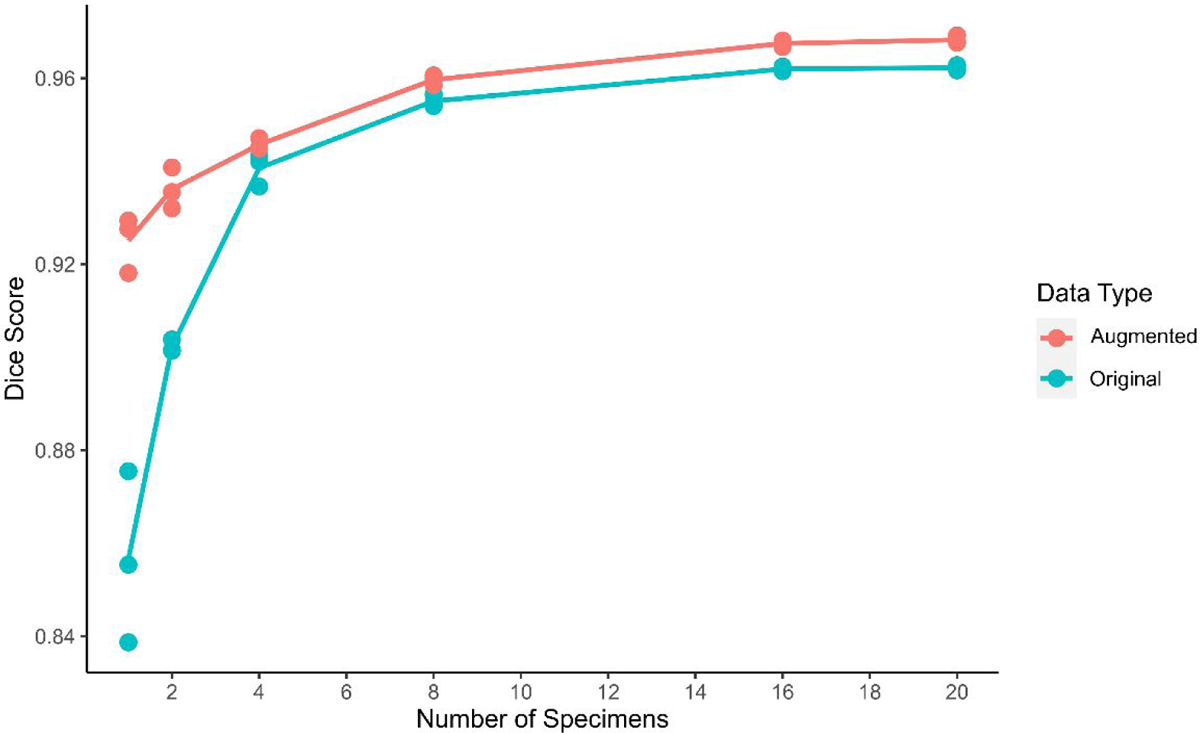
Comparison of segmentation accuracy for Biomedisa automated segmentation using average Dice scores calculated for validation data generated from 10 specimens. The plot shows increasing the training set size and implementing data augmentation improves network accuracy. All models were trained using early stopping within 25 epochs.

### 3.3 Volumetric comparison

We identified a significant overall correlation between the manually generated and network-generated total volumes for the 20 test images (F _1, 216_ = 51223, p < 0.001; figure 6), but uncovered significant differences in the degree of these correlations across training sets (F _11, 216_ = 19.660, p < 0.001). Here, the R^2^ values, serving as indicators of goodness of fit, spanned from 0.9505 for the training set with 1 specimen to 0.9998 for that containing 20 specimens with data augmentation - a difference of 0.0493.

**Figure 6.**
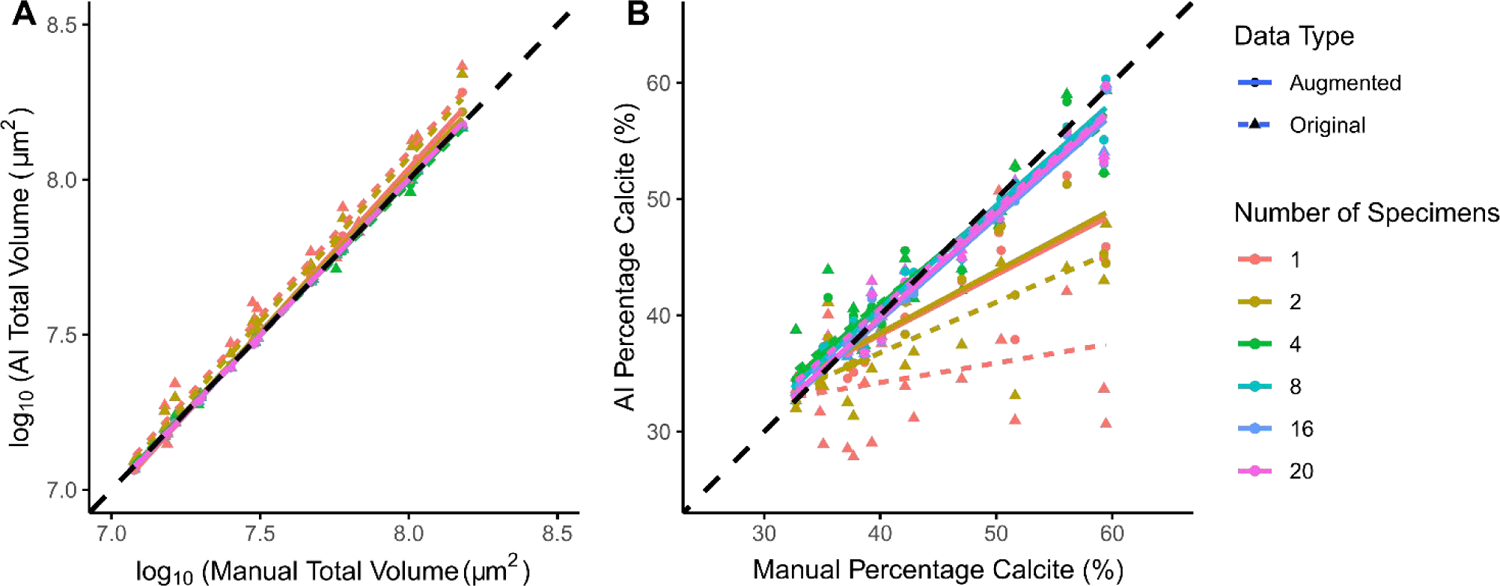
Comparison of (a) total volumes (internal and external combined) and (b) percentage calcite obtained from manual segmentation compared to those obtained using AI segmentation using 6 different training sets consisting of different numbers of training images. The predictions of total volume are generally more accurate than percentage calcite.

For percentage calcite, the correlation between the manually generated and network-generated values was also significant (F_1, 216_ = 1093.2, p < 0.001), and again significant differences were noted in the degree of correlation across training sets (F_11, 216_ = 21.367, p < 0.001). Notably, the deviation in the gradients from a perfect fit were more pronounced for the lower training for percentage calcite than for the total volumes. This difference was reflected in the R^2^ values, which ranged from 0.0608 for the training set with 1 specimen to 0.9607 for the 20-specimen set with image augmentation – a much greater difference of 0.8999.

### 3.4 Shape comparison

We expanded the analysis to compare shape estimates between the manually derived data and the six training sets for both external and internal structures. In total, 16,530 control points were generated for the external structure, while 17,325 control points were generated for the internal structure. These data points were subsequently reduced to five principal axes through non-linear Kernel principal component analysis (PCA) [23], using 1000 iterations.

Both the internal and external shape data exhibited strong and statistically significant correlations between the manually generated and network-generated shape values across all three axes, indicating a consistency in their shape estimates (figure 7; table 2). In contrast, the internal structures displayed substantial variations across the training sets as the number of images used increased from 1 to 20 across all three axes (table 2), indicating differences amongst the accuracy of different training sets.

**Figure 7.**
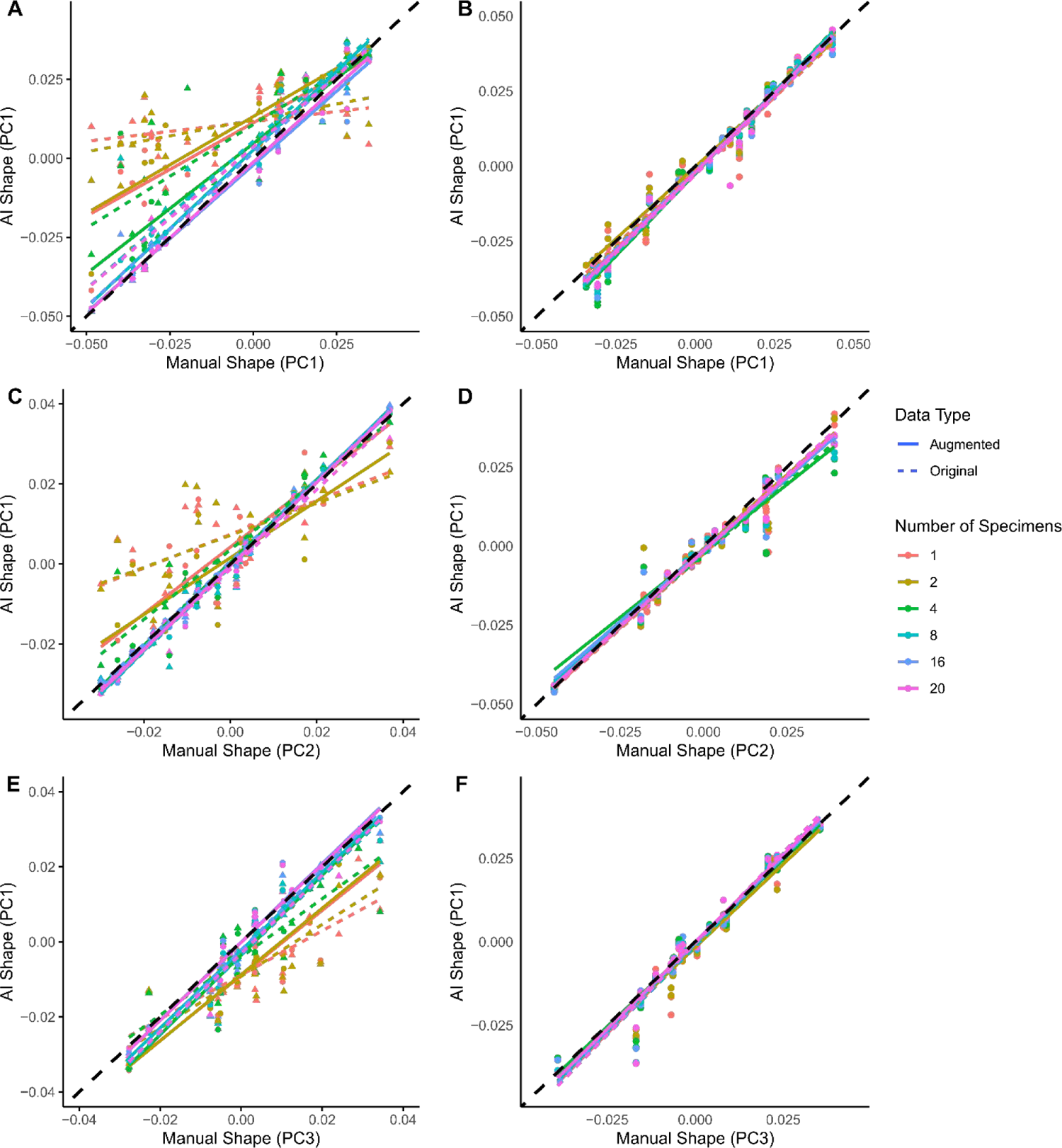
Comparison of shape for the internal (a, c and e) and external (b, d and f) across PC axes 1 (a, b), 2 (c, d) and 3 (e, f) obtained from manual segmentation compared to those obtained using AI segmentation using 6 different training sets consisting of different numbers of training images. The figure demonstrates that external calcite is easier to predict than the internal chamber space.

**Table 2.** There was no quantitative difference between manually generated and network-generated shape estimates for external and internal structures (left as F statistics on 1 and 216 degrees of freedom), but clear differences in predictive ability between external and internal shape estimates as the numbers of training images increases (right as F statistics on 11 and 216 degrees of freedom). F statistics are presented to 4 significant figures; values in bold indicate p<0.05 whereas non-bold is p>0.05.

|  | Between manual and network predictions |  | Among different numbers of training images |  |
| --- | --- | --- | --- | --- |
| Axis | External | Internal | External | Internal |
| PC1 | **7918** | **548.4** | 0.5695 | **5.442** |
| PC2 | **3474** | **1002** | 0.2491 | **3.893** |
| PC3 | **3886** | **1208** | 0.3501 | **6.997** |

## 4. Discussion

In this study, we observed a positive correlation between network accuracy and the quantity of training images, with data augmentation proving an effective tool for enhancing performance, especially for smaller training sets. External structures were extracted relatively accurately in the smaller training sets, unlike the internal structures, and size was more straightforward to extract than shape. Noting that some traits will always be more difficult to extract accurately than others, we discuss key concepts for the ongoing implementation of automated feature extraction in the biological sciences.

### 4.1 Impact of early stopping

The early stopping feature substantially reduced training time whilst effectively preserving network accuracy (see supplementary table 2). These qualities made the early stopping feature particularly valuable for larger training sets, and when applying data augmentation techniques.

Beyond improving time efficiency, the integration of early stopping features also serves to mitigate generalisation errors and overfitting [24, 25]. As a result, the application of early stopping features is widely applied when training deep learning models [26, 27]. Our findings strongly endorse the use of these early stopping features in future research using 3D image segmentation.

### 4.2 Impacts of training set size and data augmentation

Our findings reaffirm the principle that expanding the training set leads to the production of better deep learning models [28, 29], albeit with diminishing returns as accuracy approaches 100% [30]. The expansion of available training data plays a crucial role in reducing generalisation error and thus, in facilitating the achievement of optimal accuracy levels [31]. This is reflected in the stability of accuracy observed across epochs in larger training datasets, which are less affected by noise (see supplementary figures 1-12).

Importantly, the minimum number of training images to achieve accurate results and the degree to which increasing the training set size enhances network accuracy remains task-specific and context-dependent [32]. Notably, in this study, only a small number of individuals were required to achieve high accuracy scores. This achievement can be attributed to the wealth of 2D slices per specimen and the ready availability of high-quality segmentation data. Without these attributes, a larger number of individuals for training would likely be necessary, resulting in delayed increases in accuracy. Consequently, the generation of high-quality training data emerges as a factor more crucial than its sheer quantity [33], with any errors or inconsistencies within the training set likely to manifest in the deep learning models. Furthermore, the choice of the network architecture may vary in its suitability for segmenting specific materials or structures [34] meaning training sets must be selected carefully. Data augmentation is a valuable tool for enhancing network accuracy, addressing limitations stemming from insufficient training data availability. Our results strongly support that data augmentation can effectively boost model accuracy [35, 36].

Although variable by feature, we have demonstrated that it is possible to train a network with as few as one or two specimens to extract segmentation data as accurately as 8 specimens using this augmentation approach. As a result, fewer images are needed for training and thus we can significantly reduce the time spent manually segmenting. Moreover, its application in this context is straightforward and universally applicable to 3D image datasets.

### 4.3 Extracting phenotypic traits

Our results suggest extracting the internal structure of planktonic foraminifera poses a greater challenge compared to extraction of the external structure, evidenced in the measures of percentage calcite and the shape of the internal structure (see figures 5b, 6). This discrepancy in difficulty can be likely be attributed to material homogeneity, which is reflected in low contrast differences within CT scans [37, 38]. This challenge is particularly prominent in the case of planktonic foraminifera, where the internal structure contains sedimentary infill and nannofossil ooze with densities similar to external calcite [39, 40]. The quantity of this infill is likely to vary among specimens, ranging from abundant to absent, while the presence of external calcite remains consistently detectable. Consequently, to alleviate the impact of irrelevant features, such as sediment infill, on trait extraction, we need training sets that encompass a broader array of images, representing a fuller sample of the entire image population compared to datasets without these elements. This can be achieved by either increasing the size of the training set or employing data augmentation techniques, thereby facilitating more accurate predictions.

Additionally, further enhancements in the accuracy of selected networks and mitigations of irrelevant features can be achieved through the application of post-processing tools such as the removal of connected components and the application of smoothing methods to eliminate noise from deep learning outputs [41, 42] and hold the potential to yield more precise segmentation data.

### 4.4 Implications for future studies

There is a growing demand in the field of phenomics for the integration of machine learning techniques to perform image segmentation, driven by the increasing abundance of available data [43]. Unlike manual segmentation, which is inherently prone to observer bias [44], deep learning methods offer a robust means to ensure greater consistency across measurements [45], facilitating efficient and scalable comparative studies. The complexity and size of phylogenies [46] and birth-death models [47] are only getting larger, with the use of continuous traits [48] and geometric morphometric approaches [49] also now commonplace. Thus, there is a need to develop morphometric approaches that can keep pace with these advances.

The application of the automated segmentation and the data augmentation techniques applied here can be used universally to improve data acquisition. They integrate seamlessly with widely used segmentation software like Dragonfly, Avizo/Amira, ImageJ, and MITK, so can be incorporated into existing workflows. They also accommodate various input sources, encompassing images obtained through X-ray synchrotron microscopy [50], soft microscopy [51], and magnetic resonance imaging (MRI) [52]. This versatility empowers the workflow to conduct large-scale analyses efficiently and with a high level of accuracy within reasonable timeframes. With the provision of suitable input data [53], these trained networks become accessible to anyone for segmentation, regardless of their expertise in the morphology of the selected specimen.

The ability to extract different 3D features using deep learning remains likely to be context specific, with a variety of features such as the presence of artefacts and irregularities [54], broken or incomplete specimens [55], and low contrast images [42] each requiring individual solutions to overcome. Improvement in the quality of image data acquisition and selection of specimens can aid in mitigating some of these factors [6], however, these must coincide with good training sets.

The degree of tolerable error is also influenced by the quantification required. Here, our results demonstrate the extracting the total volume was easier than percentage calcite and shape. The variation in total volume is much greater and is less complicated to measure, thus is less confounded by inaccurate measurements compared to its counterparts. The landmark-free method applied here, for instance, requires the generation of meshes, and scaling and alignment of the data which further produces error. Researchers should thus consider their measurements when training their neural networks.

## Conclusion

We provide compelling evidence that leveraging deep learning for automated segmentation not only yields equivalent results to manual segmentation but also drastically reduces the time required for extracting 3D features from images. Data augmentation emerged as a powerful tool, elevating network accuracy and enhancing the efficacy of smaller training sets. However, it is imperative to acknowledge that the accuracy of feature extraction is contingent on the specific traits targeted. The observed differences for internal and external traits extracted underscores the crucial need for thoughtful consideration when training deep learning networks for specific applications.

## Data accessibility

All computer code and final raw data used in the paper is available at https://github.com/JamesMulqueeney/Automated-3D-Feature-Extraction. All other data is available in the Dryad Repository: DOI: 10.5061/dryad.x69p8czrb.

## Authors’ contributions

J.M.M, T.H.G.E and A.G designed the concept of the study. A.B performed all the taxonomic classifications and picking of the foraminiferal specimens. A.S.B performed the computer tomography scanning of all specimens. J.M.M, A.S.B and M.S performed manual image segmentation across the 50 specimens. J.M.M trained the AI models and performed the volumetric and shape comparisons. J.M.M wrote the initial draft manuscript, which was iterated with A.G. and T.H.G.E. All authors provided ideas and discussion to the paper and have read and approved the final version.

## Acknowledgments

The authors would like to thank Philipp D. Lösel whose advice on performing the AI network training was invaluable, and all the other members of the Goswami and Ezard Labs who generously provided ideas for the concept of the paper.

